# An adversarial scheme for integrating multi-modal data on protein function

**DOI:** 10.1101/2025.01.16.633332

**Authors:** Rami Nasser, Leah V Schaffer, Trey Ideker, Roded Sharan

## Abstract

In order to begin to decipher the structure of the cell, we need to integrate multiple types of data of different scales on subcellular organization. Such integration requires dealing with multiple data modalities and with missing data. To this end, we developed MIRAGE, a multi-modal generative model for integrating protein sequence, protein-protein interaction and protein localization data. Our approach successfully learns a joint embedding space that captures the complex relationships between these diverse modalities. We evaluate our model’s performance against existing methods, obtaining superior performance in several key tasks, including protein function prediction and module detection. MIRAGE source code is available at https://github.com/raminass/MIRAGE.

## 1 Introduction

Sparsity and incomplete data present significant challenges in bioinformatics, hindering the analysis and interpretation of large biological datasets [23]. These issues necessitate the development of specialized computational algorithms and data imputation methods to accurately predict missing values and extract meaningful insights from incomplete biological information. A single protein encompasses multiple biological dimensions: its amino acid sequence reveals insights into its structure and molecular function; its protein-protein interactions (PPIs) reflect the biological processes it takes place in and provide information on its subcellular organization; and its localization images illustrate its spatial distribution within cellular compartments. This multifaceted nature of proteins provides various sources of information for learning meaningful representations by integrating different biological modalities. Ideally, a unified joint embedding space would allow for an integrated representation of proteins by aligning these diverse modalities.

However, acquiring comprehensive datasets that include all these modalities is often impractical due to the high costs and complexities associated with experimental data collection about the interactions and localization of a protein. Thus, state-of-the-art integration methods consider only a subset of these modalities. For instance, some approaches focus on integrating localization images with PPI information [28], while others aim to connect sequence data with localization images [6]. Similar approaches have been successful in other fields, such as combining images with text or audio in computer vision [1, 3]. Nevertheless, these methods typically operate on limited pairs of modalities, resulting in embeddings that are restricted to the specific combinations used during training. A recent study [12] attempted to address this limitation by using images as a central point to connect with other types of data. While this is a step forward, it depends on comprehensive image data which is not available as of yet in the protein world.

Our work addresses this gap by proposing the MIRAGE (Multi-modal Integrative Representation using Adversarial Generative Embedding) model that learns a joint embedding space across the three aforementioned modalities: sequence, interaction and localization. Importantly, our model does not require full information, allowing us to represent proteins for which information on one or two modalities is missing. Our approach draws inspiration from CycleGAN [35], adapting its concept of bidirectional translation to the domain of biological data modalities. In our model, different modalities are encoded into a shared latent space, from which we can generate other modalities. This creates a cycle of translations: modality A can be used to generate modality B, and the generated B can be used to reconstruct A, ensuring consistency and information preservation across modalities. This methodology enables the translation and generation of one modality from another, offering a novel solution to the pervasive issue of data scarcity and incompleteness in biological research.

We demonstrate the effectiveness of our multi-modal generative model by integrating protein sequence data, protein-protein interaction information, and subcellular localization images to construct a hierarchical map of subcellular organization. Our results show that our approach successfully learns a joint embedding space that captures the complex relationships between these diverse modalities. We evaluate our model’s performance against existing methods for protein representation learning, including those that focus on a single modality or on pairs of modalities. Our framework demonstrates superior performance in several key tasks, including protein function prediction, module detection and data generation for missing modalities.

## 2 Methods

### 2.1 Problem formulation and model outline

Let *X* = {*X*_1_, *X*_2_, ߪ;, *X*_*M*_} represent the set of *M* different modalities for protein representation (e.g., sequence (SEQ), interaction (PPI) and localization (IMG)), where 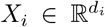. The goal is to embed 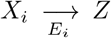 into a shared laent space *Z* ∈ ℝ^*l*^, and to generate 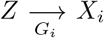 modality vector from any latent modality *j*, where *E*_*i*_ and *G*_*i*_ are parameterized mapping functions.

We employ adversarial learning, where discriminators *D*_*i*_ are trained to distinguish between real and generated samples of each modality. The key innovation lies in feeding unaligned modality embeddings to the framework, allowing the model to learn cross-modal relationships without requiring full *M*-tuple data from the same protein. Formally, for any pair of modalities (*i, j*), we can perform translations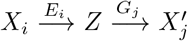, where 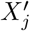 is the generated version of modality *j* from modality *i*. Our approach, MIRAGE, is illustrated in Figure 1. Importantly, MIRAGE’s translation process is independent of the specific proteins used in the input modality, enabling flexible cross-modal generation even when data for all modalities is not available for every protein. This overcomes the requirement for aligned data that often drastically reduces the volume of available training samples as demonstrated in Supplementary Figure B.3.

**Fig. 1:**
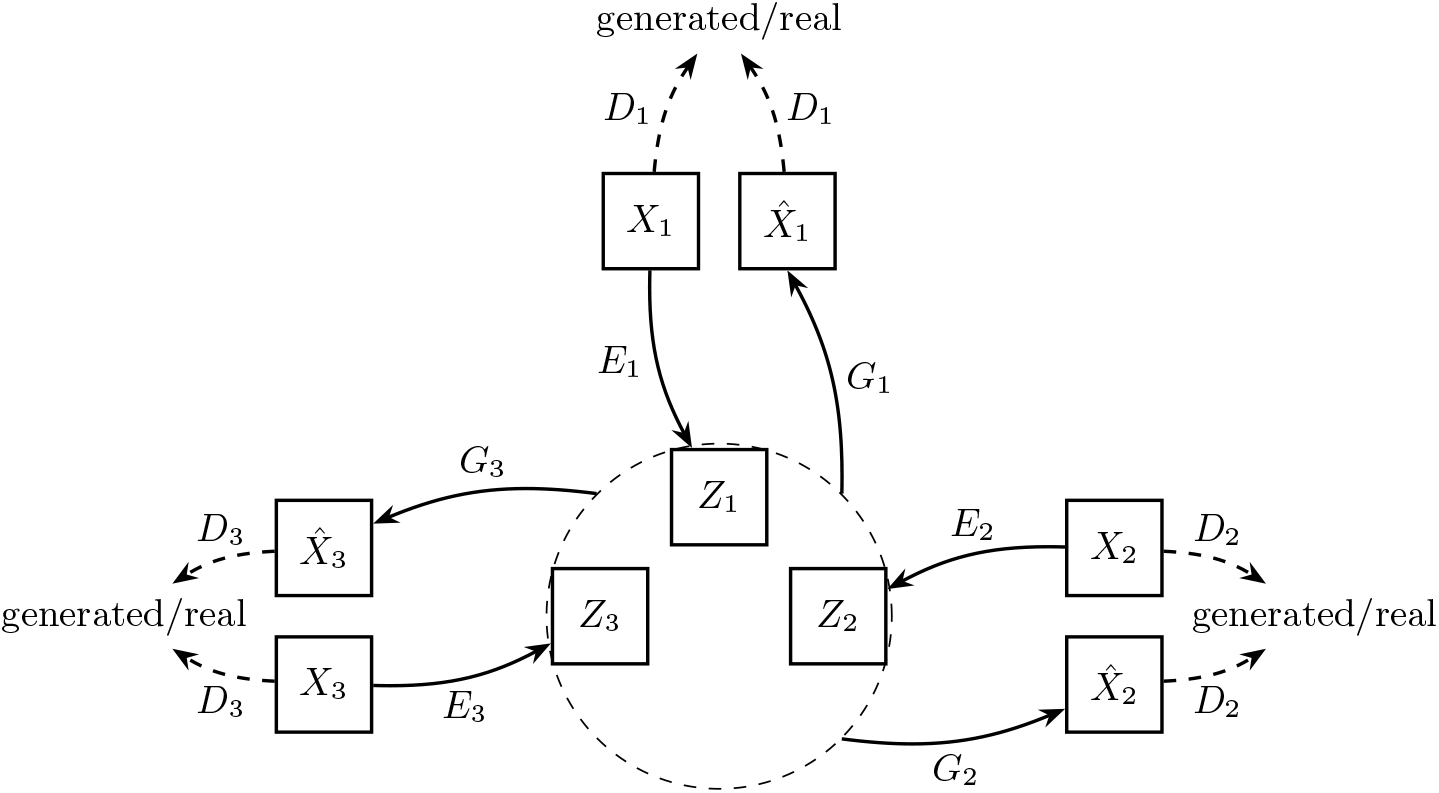
The MIRAGE scheme. For each modality it learns a mapping *E* : *X* → *Z* to latent space (dashed circle) and a generator from latent space to modality *G* : *Z* → *X*. Real and generated samples are classified using a parameterized discriminator 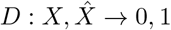.

Our objective function consists of three main components, which are described in detail in the Supplementary Methods: (i) adversarial loss ℒ_*gan*_, which ensure that the distribution of the generated modality *i* matches the data distribution in the target domain; (ii) reconstruction loss ℒ_*rec*_, which enforce consistency between the latent space representation and the original domain; and (iii) latent-cycle-consistency loss ℒ_*cyc*_, introduced to prevent contradictions between the learned mappings *E* and *G*, ensuring alignment in the latent space for the same protein encoded from different modalities. The full loss function is a weighted sum of these loss terms:

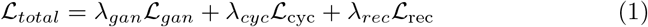

where *λ*_*gan*_, *λ*_*cyc*_, *λ*_*rec*_ control the relative importance of each loss term. Computationally, this setup requires learning 2×*M* parameters, making the complexity linear with respect to the number of modalities, in contrast to a direct mapping between all modalities, which would lead to quadratic complexity.

Full details on the model architecture and its training appear in the Supplementary Methods.

### 2.2 Application to HEK239T

We applied MIRAGE to data coming from three different modalities measured in HEK293T cells. Specifically, we integrated protein sequence data from UniProt for 20,218 human proteins, alongside immunofluorescence images from the Human Protein Atlas [32] and protein-protein interaction data from the Bioplex network [19]. The protein interaction network contains 14,032 proteins and 127,732 interactions, while the image dataset includes 1,125 immunofluorescence images. Dataset sizes and overlaps are depicted in Supplementary Figure B.3.

After learning, in order to construct a representative embedding for each protein that accounts for its image-based, network-based and sequence-based embeddings, we fuse those embeddings into a joint representation. While there are many potential fusion functions [24], we simply use concatenation: *Z* = [*Z*_1_|*Z*_2_|*Z*_3_].

In recent works, there has been a growing shift toward using embedded or encoded features instead of raw data, enabling models to extract rich, abstract representations that capture essential aspects of the data [7]. By breaking large tasks into smaller, manageable ones, intermediate representations can be learned and then leveraged for the main objective. For example, in ESM-2 [25], masked language modeling (MLM) was first applied to learn sequence embeddings, which were subsequently used to train a 3D structure prediction model, in contrast to end-to-end approaches like AlphaFold [20]. Another example is in computer vision, where image representation is learned for downstream tasks such as: classification, segmentation [10, 31]. Following this practice, MIRAGE also utilizes embeddings as input, incorporating multiple modalities in a joint embedding space rather than working directly with raw data. This approach enhances flexibility and improves performance in complex tasks.

For sequence encoding, we utilized the ESM-2 [25] model to obtain sequence embeddings from the raw sequence data. For image encoding, we use DenseNet [30]. For network encoding, we employed node2vec [16]. These encodings allow for direct comparison with previous works that have used similar methods [6].

### 2.3 Performance evaluation

To assess the performance of our method, we utilized the recently developed BIONIC benchmark [11] with the following tasks: (i) protein module detection and (ii) supervised protein function prediction. While BIONIC originally employed hierarchical clustering for module evaluation, we opted for the state-of-the-art Louvain clustering algorithm [8] due to its computational efficiency and effectiveness in identifying community structures in large networks. Our evaluation was conducted using human protein module benchmarks derived from multiple well-established biological databases. These included KEGG pathways [21], excluding metabolic pathways to focus on signaling and regulatory modules; Gene Ontology (GO) annotations [4], specifically targeting Cellular Components (CC) and Biological Processes (BP); and CORUM complexes [13], which provide a curated set of mammalian protein complexes. This diverse set of benchmarks allowed us to evaluate our method’s performance across various biological contexts and scales, ranging from specific protein complexes to broader functional modules and pathways.

## 3 Results

We applied MIRAGE to integrate sequence, interaction and localization information in HEK293T cells to construct a hierarchical map of protein subcellular organization. As a sanity check, we compared the MIRAGE joint embedding to those obtained from each modality separately. As illustrated in Figure 2, our method consistently outperformed single-modality approaches across all benchmark datasets. The joint embeddings produced by MIRAGE better align with known biological structures compared to those derived from any individual modality. This superior performance underscores the effectiveness of our approach in capturing complementary information from diverse data types, resulting in more biologically relevant and informative representations.

**Fig. 2:**
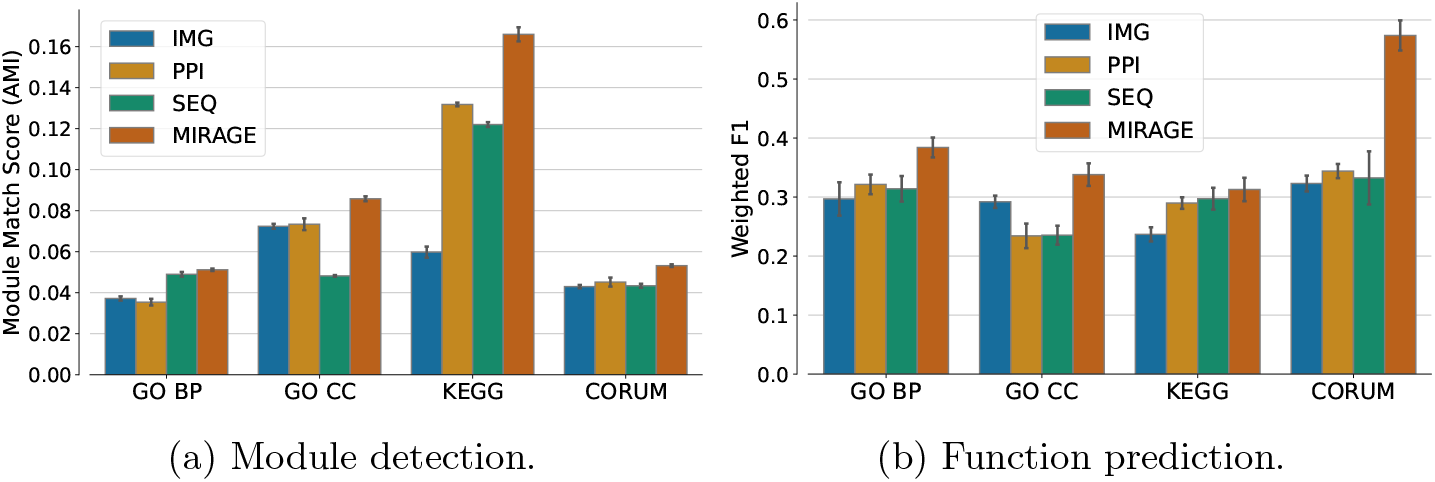
Performance evaluation of the learned joint representation against the individual modalities. A) Module detection performance using the adjusted mutual information (AMI) measure. B) Function prediction performance evaluated by Weighted F1 which is the average per-class F1 score.

While MIRAGE seamlessly integrates all three input modalities, methods such as MUSE [6] and DICE [28] are constrained to integrating only two modalities at a time. To compare to those methods, we performed three distinct experiments, each combining a pair out of the three modalities. For each modality pair, we applied the same evaluation protocol described earlier. The results of these pairwise integration experiments are presented in Figure 3. Remarkably, MIRAGE consistently outperformed the benchmark methods across almost all modality combinations and evaluation benchmarks.

**Fig. 3:**
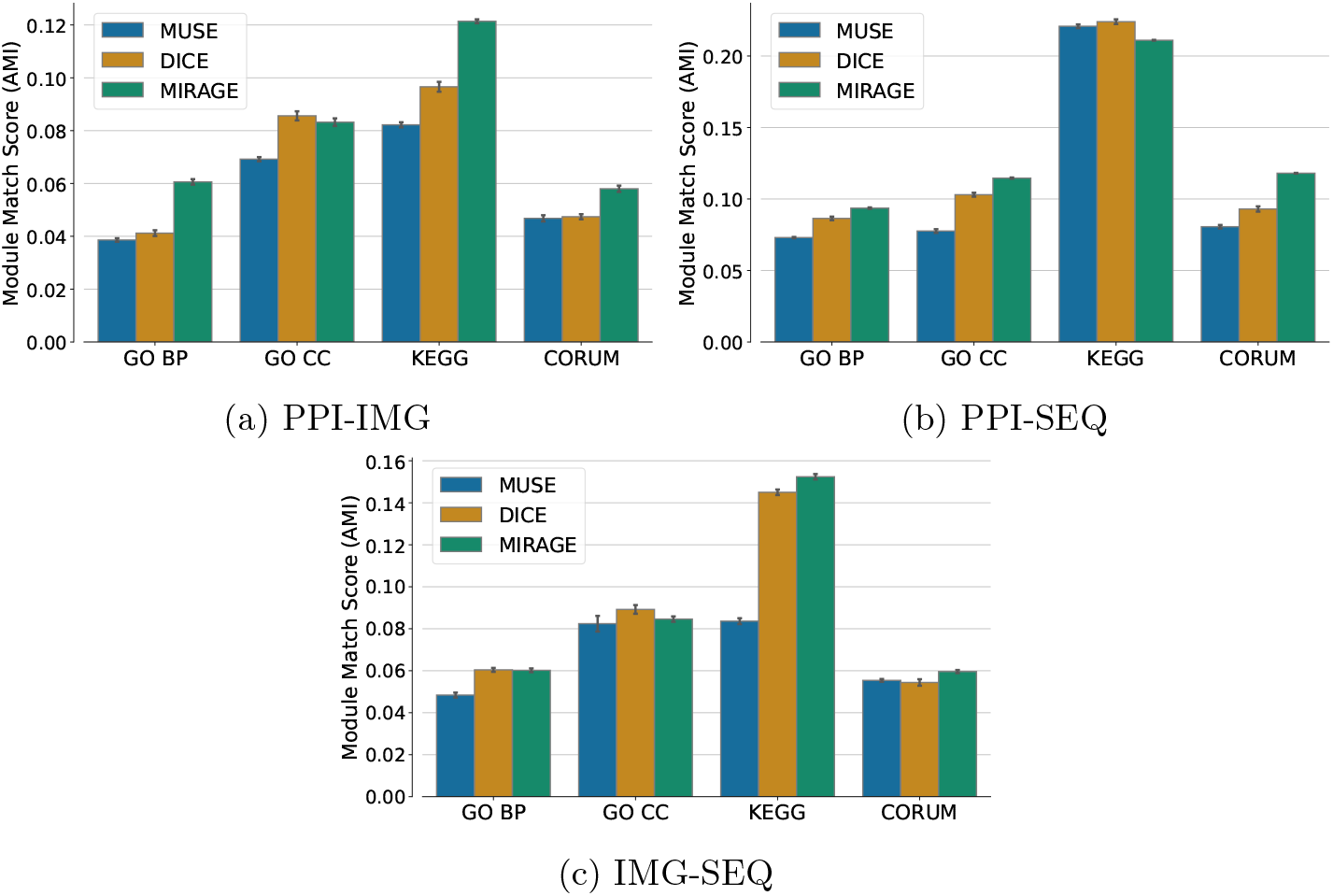
Performance evaluation of the learned joint representation using MIRAGE against previous methods for integrating pairs of modalities (MUSE and DICE).

To comprehensively evaluate MIRAGE in the context of integrating three or more modalities, we conducted a comparative analysis against alternative approaches capable of handling multi-modal data. Due to the scarcity of well-established models designed to operate on three or more modalities simultaneously, we selected two baseline methods: simple feature concatenation (CONCAT) and a multi-modal autoencoder (MMAE) which is considered a strong baseline for multi-modal integration [5]. The CONCAT method serves as a naive baseline, directly combining features from different modalities without learning inter-modal relationships. In contrast, the MMAE represents a more sophisticated approach, combining multi-modal data through a bottleneck layer that can reconstruct original features, a technique that has shown promise in various multi-modal learning tasks [29]. The results of this comparison, constrained to proteins with data in all modalities, are presented in Figure 4. Remarkably, MIRAGE consistently outperformed both CONCAT and MMAE across all benchmarks and evaluation metrics.

**Fig. 4:**
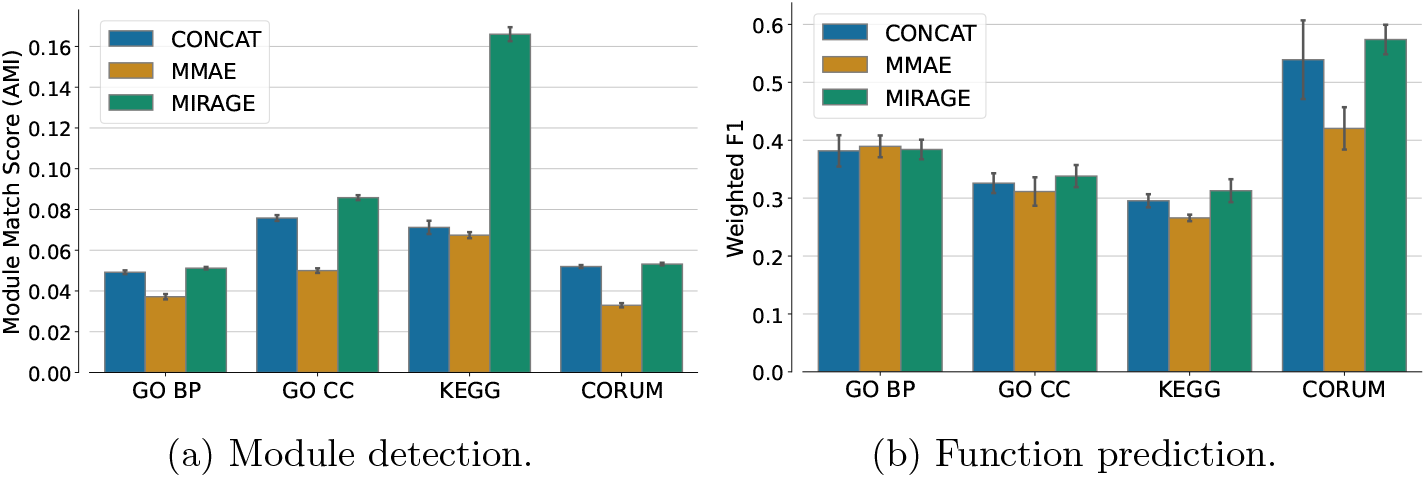
Performance evaluation of the learned joint representation against other multi-modal methods (CONCAT and MMAE).

**Fig. 5:**
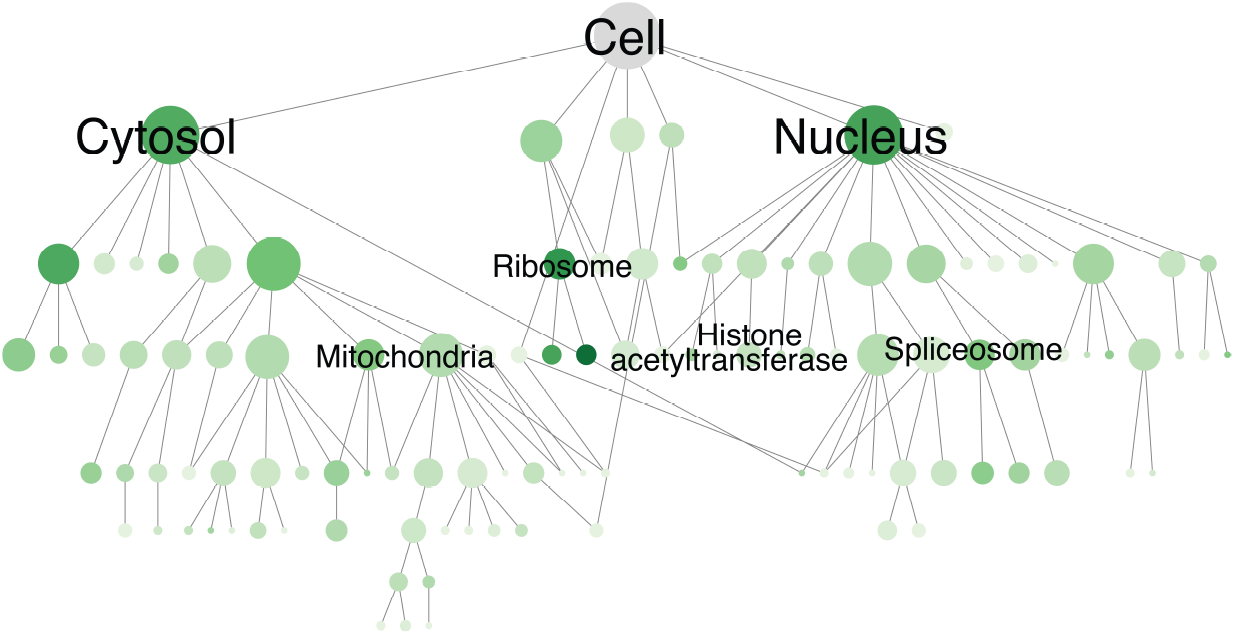
Hierarchy of protein assemblies constructed using the MIRAGE embeddings. Nodes represent protein assemblies and edges represent hierarchical containment. Node size is proportional to number of proteins. Nodes are shaded based on overlap with known cellular components.

### 3.1 Hierarchical cell map

After establishing the utility of MIRAGE, we applied it to construct a hierarchical map of cell structure. To this end, we clustered the proteins according to the similarities between their integrative embeddings at multiple resolutions [34]. For comparison purposes, we focused on the 907 proteins present in all three modalities. The hierarchy contains 111 clusters, including 62 clusters that overlap significantly with a component in the Gene Ontology, CORUM or HPA (hypergeometric test, FDR *<*10% and Jaccard index *>*10%). We recovered assemblies across scales, including large compartments (e.g., nucleus and cytosol) and small compartments (e.g., histone acetyltransferase complex and ribosomal complex). In comparison, a previous integration of interaction and image information using DICE [28] identified only 46 clusters that overlapped with known components at the same thresholds. This comparison suggests that MIRAGE embeddings better capture known biological systems.

### 3.2 Embedding Alignment and generating missing information

Despite MIRAGE’s foundation in generative adversarial networks (GANs), we observed a notable alignment property in MIRAGE’s joint embeddings. This phenomenon is reminiscent of the characteristics typically associated with contrastive learning approaches, where matching pairs are drawn closer together in the embedding space while non-matching pairs are more uniformly distributed [33]. To investigate the alignment properties of our joint embedding, we employed the UMAP dimensionality reduction algorithm [27] for visualization. The results, presented in Figure 6, reveal a striking alignment among the three modalities for each protein. This observed alignment is particularly noteworthy given that MIRAGE does not explicitly optimize for this property. The emergence of such alignment suggests that our model has successfully captured the intrinsic relationships between different modalities of the same protein, effectively learning a shared representation space.

**Fig. 6:**
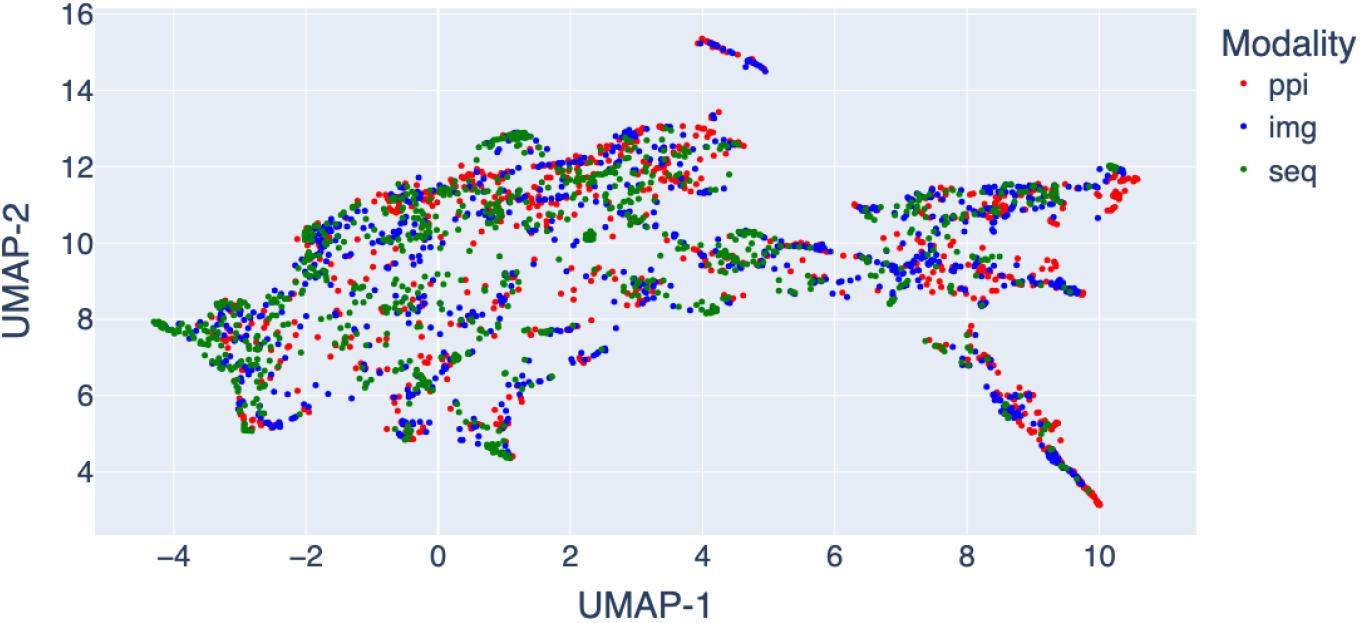
UMAP visualization of latent MIRAGE embeddings.

As image information is the scarcest, we evaluated our model’s ability to generate it using data from the other two modalities. To assess the quality of these generated image embeddings, we employed the Fréchet Inception Distance (FID) metric [18] (Supplementary Methods). We compared the FID scores of our generated images embeddings against those produced by a K-Nearest Neighbors (KNN) approach, using the true image embeddings as a reference. Our model consistently achieved lower FID scores compared to KNN, indicating superior performance (see Figure 7). This shows the effectiveness of our method in generating high-quality protein image representations from PPI, sequence or a combination of both (Supplementary Methods). These results underscore the potential of our model to fill gaps in protein datasets, which could have significant implications for downstream tasks.

**Fig. 7:**
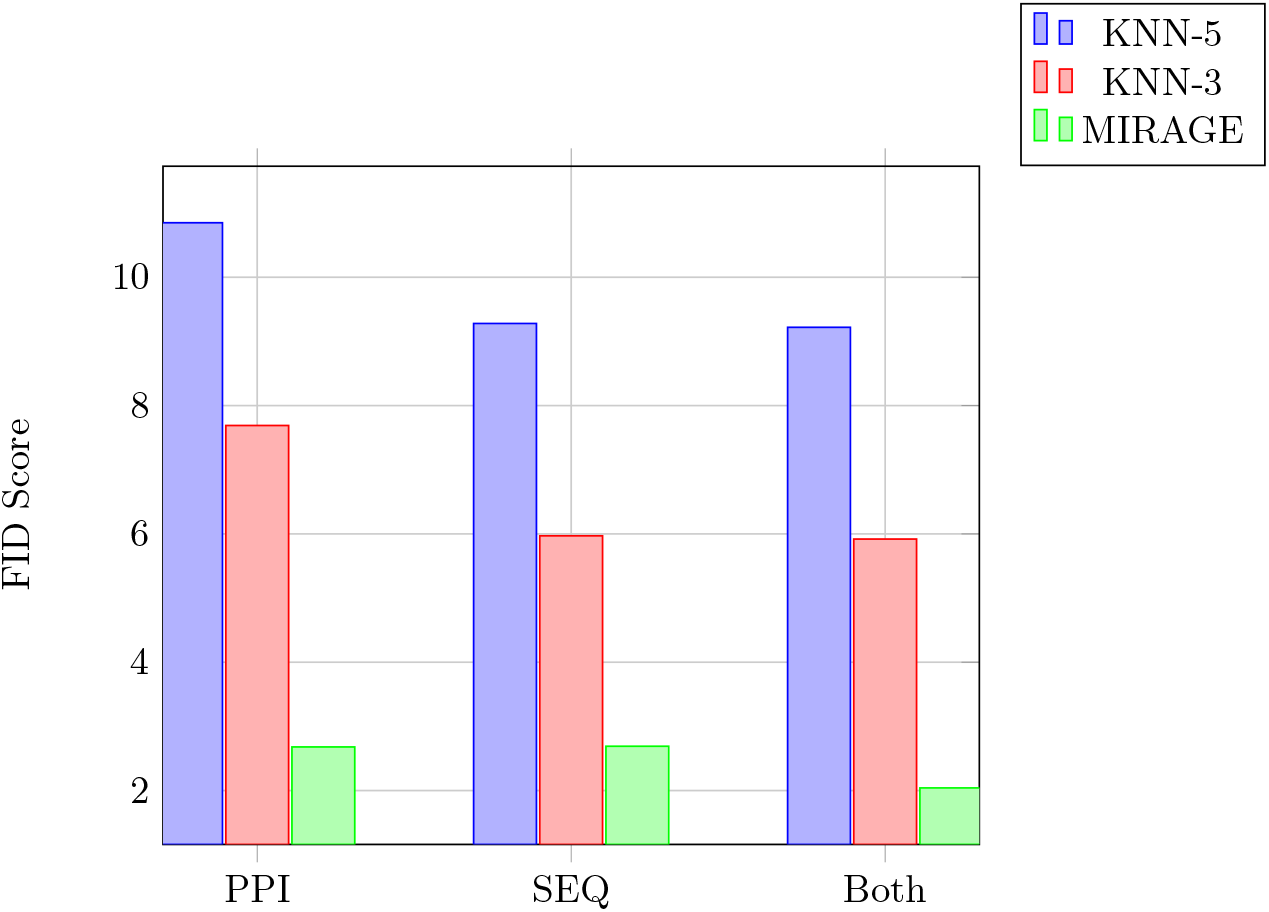
FID scores for generated images using PPI information, sequence data, or both. The performance of MIRAGE is compared to k-NN with *k* = 3 or *k* = 5.

## Conclusions

MIRAGE is a multi-modal generative model for integrating protein sequence, protein-protein interaction and protein localization data for constructing a hierarchical map of subcellular organization. Our approach successfully learns a joint embedding space that captures the complex relationships between these diverse modalities. By enabling the use of unaligned data, our model can exploit a broader range of information, potentially leading to more robust and generalizable representations. This unaligned training paradigm offers several advantages. First, it substantially increases the quantity of data that can be utilized for training, as it removes the constraint of finding perfectly matched samples across modalities. Second, it enhances the model’s ability to learn more flexible and diverse mappings between modalities, potentially capturing a wider range of inter-modal relationships.

## Acknowledgments

RN was supported by a fellowship from the Edmond J. Safra Center for Bioinformatics at Tel-Aviv University. RS was supported by a research grant from the Israel Science Foundation (grant no. 1692/24).

## A Supplementary Methods

### Adversarial loss

Generative Adversarial Networks (GANs) [15] are a class of deep learning models consisting of two neural networks, a generator and a discriminator, trained simultaneously through adversarial learning. In traditional GANs, the discriminator is typically trained using binary cross-entropy loss to distinguish between real and generated samples, while the generator aims to produce samples that can fool the discriminator. However, this approach can lead to training instability and vanishing gradients. To address these limitations, we adopted the Least Squares Generative Adversarial Network (LSGAN) [26] approach in our work. LSGAN replace the cross-entropy loss function in the discriminator with a least squares loss. This modification provides several advantages: (i) it helps mitigate the vanishing gradients problem often encountered in GAN training; (ii) and it addresses the issue of the discriminator learning too quickly and overpowering the generator, which can lead to training instability. This results in improved stability during the training process compared to regular GANs.

For the discriminator the loss is:

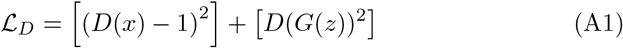

where 1 represents the target label for real samples, and only the discriminator *D* is updated during this step.

For the generator the loss is:

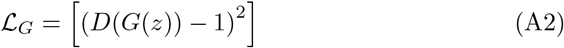

where 1 represents the target label for generated samples to fool the discriminator, and only the generator *G* is updated during this step.

We further incorporate the gradient penalty [17] approach, which enhances the stability of GAN training by regulating how quickly the discriminator’s output can change with respect to its input. Specifically, the gradient penalty adds a term to the loss function that discourages the discriminator from making overly confident predictions based on small changes in input. This helps to prevent the discriminator from becoming too powerful too quickly, which can lead to training instability. The gradient penalty is added to the original ℒ_*D*_ loss, resulting in the following adversarial objective:

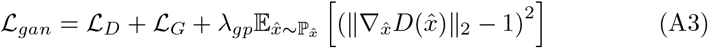

where *λ*_*gp*_ is a constant that controls the strength of the gradient penalty [5], 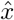 is the input sampled along the line between real and generated samples, sampled only from real,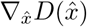 is the gradient of the discriminator with respect to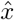, ∥ · ∥ _2_ is the L2 norm, and the term 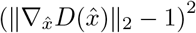 encourages the gradient norm to be close to 1.

### Cycle Consistency Loss

The cycle consistency loss, originally introduced for CycleGAN [35], helps ensure that the translation process is bijective and preserves important features of the input. By extending this concept to the latent space, we aim to enforce consistency in the learned representations (Supplmentary Figure B.2), which can lead to more stable and meaningful translations. This latent cycle consistency loss, allows many to many mapping without fixing pairs to perform a full cycle as in [35], also it contribute to better alignment of translations in the latent space. Formally, the loss is:

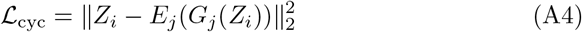

where *Z*_*i*_ = *E*_*i*_(*X*_*i*_) is the latent representation of input modality *i*. We randomly select target modality *j* and generate to that modality, then encode back to latent space(Supplementary Figure B.2).

### Reconstruction Loss

In addition to the cycle consistency loss, we utilize a reconstruction loss from the latent space back to the original space. This additional constraint helps to preserve important features of the input during the translation process and encourages the model to learn a meaningful and diverse mapping between domains. By enforcing this reconstruction, we aim to avoid the problem of mode collapse, by generator produces a limited variety of outputs regardless of the input. This approach is inspired by the original CycleGAN paper [35], and further explored in the augmented CycleGAN model [2]. We use the following reconstruction loss:

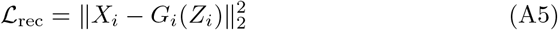

where *Z*_*i*_ = *E*_*i*_(*X*_*i*_) is the latent representation of modality *i* (Supplementary Figure B.1).

### Fréchet Inception Distance (FID)

The Fréchet Inception Distance (FID) score between two distributions *P*_*r*_ (real data) and *P*_*g*_ (generated data) is given by:

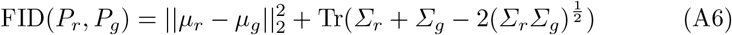

where *µ*_*r*_ and Σ_*r*_ are the mean and covariance of the real data features, and *µ*_*g*_ and Σ_*g*_ are the mean and covariance of the generated data features.

FID is particularly suitable for this task as it captures both the quality and diversity of generated images, providing a comprehensive measure of how well the generated distributions match the real data distribution [9].

### Implementation details

We implemented MIRAGE using PyTorch and conducted experiments on a Linux machine equipped with an NVIDIA TITAN Xp GPU. The source code is available at https://github.com/raminass/MIRAGE. Our network architecture utilizes a latent dimension of 128 and a hidden dimension of 512. We initialized the model weights using Xavier initialization [14]. We employed a batch size of 32 for all experiments, as it provided optimal stability for adversarial training. Smaller batch sizes such as 8 were explored but resulted in unstable training dynamics, as illustrated in Figure B.5. For optimization, We employed the Adam optimizer [22] with a learning rate of 0.0002 and *β*_1_ = 0.5, following established practices in GAN training [31]. Our training schedule maintained a constant learning rate for the first 100 epochs, followed by a linear decay to zero over the subsequent 100 epochs, a strategy that has been shown to enhance stability and convergence in GAN training [35]. Notably, we update the discriminator using only the samples produced by the most recent generator iteration. This approach was adopted after observing that using older or buffered samples led to an overpowered discriminator, potentially destabilizing the training process see Figure B.4. In our loss function, we employed multiple components weighted by specific hyperparameters to balance their contributions. The total loss is computed as a weighted sum of the GAN loss, cycle consistency loss, reconstruction loss, and gradient penalty, specifically we use the following weighting parameters same as in [35]: *λ*_*gan*_ = 1, *λ*_*cyc*_ = 10, *λ*_*rec*_ = 10, *λ*_*gp*_ = 1.

## B Supplementary Figures

**Fig. B.1:**
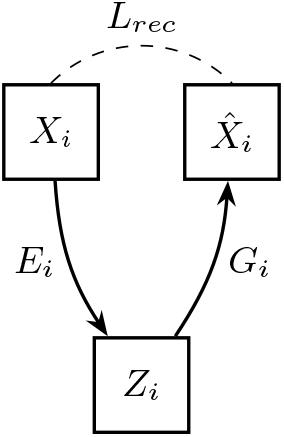
Short cycle loss (*L*_*rec*_) *X*_*i*_ ≈ *G*_*i*_(*E*_*i*_(*X*_*i*_)).

**Fig. B.2:**
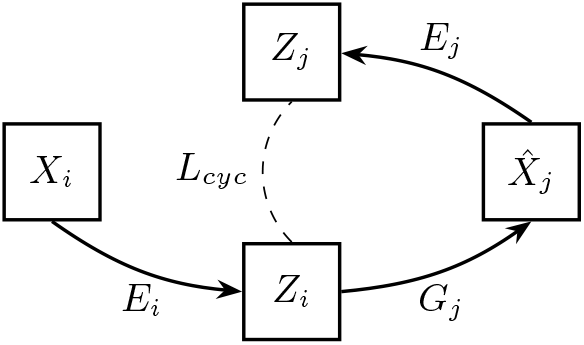
cycle consistency loss in the latent space (*L*_*cyc*_) when we translate from 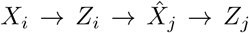 and constrain *Z*_*i*_ ≈ *Z*_*j*_, target modality *j* is randomly chosen.

**Fig. B.3:**
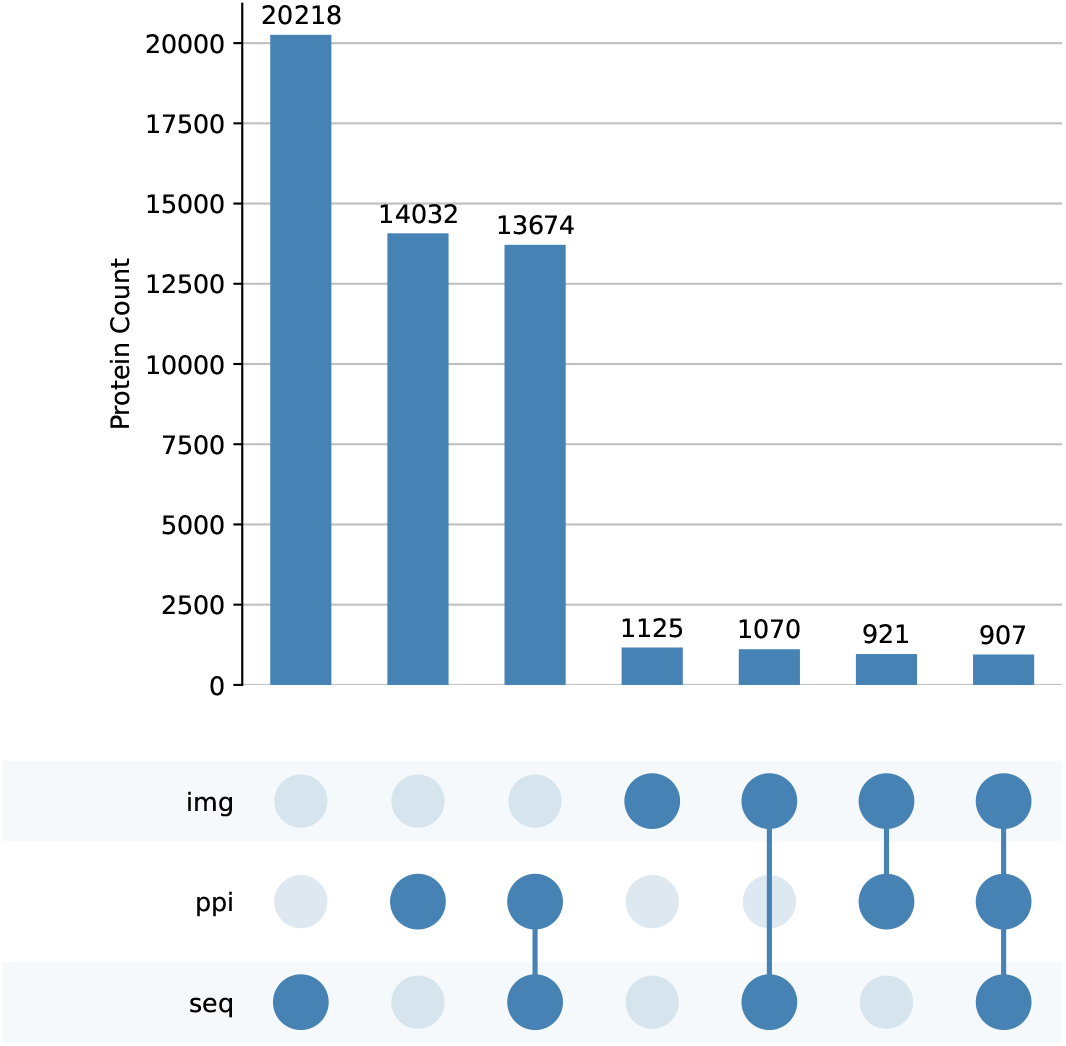
Sizes of datasets and their intersections.

**Fig. B.4:**
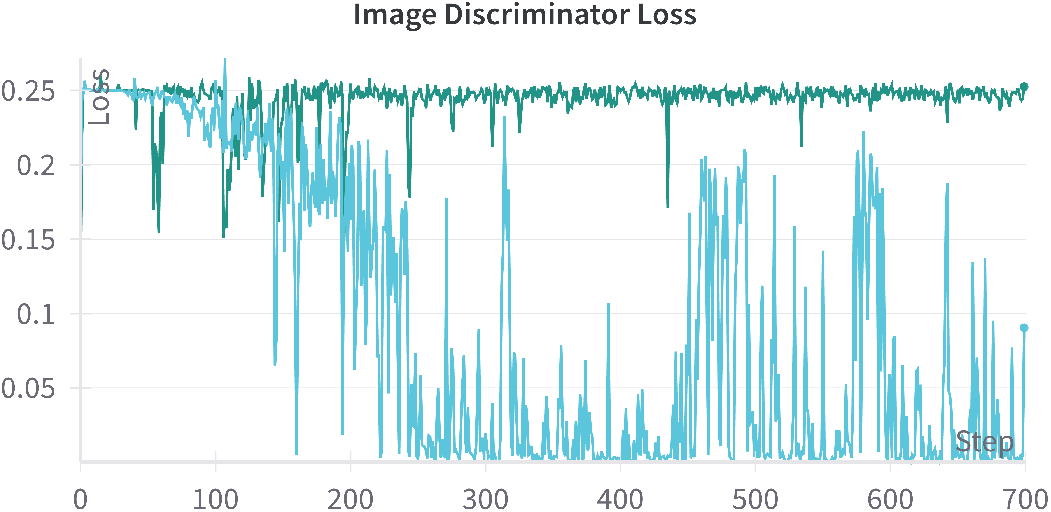
Discriminator loss during training with and without image buffering. The blue line represents the loss when employing an image buffer, while the green line shows the loss without buffering.

**Fig. B.5:**
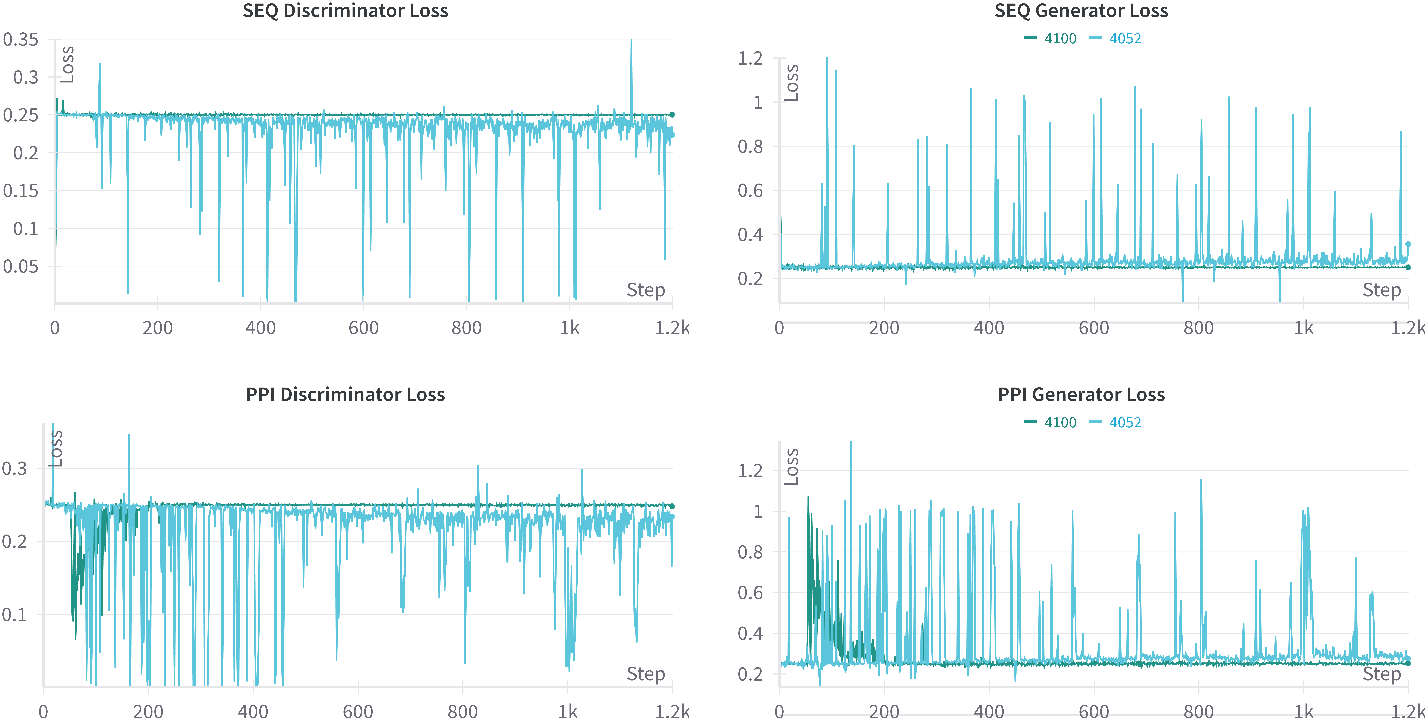
Discriminator and generator loss during training with different batch sizes. The green line represents the loss when using a batch size of 32, while the blue line indicates the loss with a smaller batch size of 8. The results demonstrate that the larger batch size of 32 leads to more stable adversarial training, whereas the smaller batch size of 8 results in increased instability, as evidenced by the fluctuations in loss over training epochs.

## References

1. J.-B. Alayrac, J. Donahue, P. Luc, A. Miech, I. Barr, Y. Hasson, K. Lenc, A. Mensch, K. Millican, M. Reynolds, et al. Flamingo: a visual language model for few-shot learning. Advances in neural information processing systems, 35:23716–23736, 2022.

2. A. Almahairi, S. Rajeshwar, A. Sordoni, P. Bachman, and A. Courville. Augmented cyclegan: Learning many-to-many mappings from unpaired data. In International conference on machine learning, pages 195–204. PMLR, 2018.

3. R. Arandjelovic and A. Zisserman. Look, listen and learn. In Proceedings of the IEEE international conference on computer vision, pages 609–617, 2017.

4. M. Ashburner, C. A. Ball, J. A. Blake, D. Botstein, H. Butler, J. M. Cherry, A. P. Davis, K. Dolinski, S. S. Dwight, J. T. Eppig, et al. Gene ontology: tool for the unification of biology. Nature genetics, 25(1):25–29, 2000.

5. T. Baltrušaitis, C. Ahuja, and L.-P. Morency. Multimodal machine learning: A survey and taxonomy. IEEE transactions on pattern analysis and machine intelligence, 41(2):423–443, 2018.

6. F. Bao, Y. Deng, S. Wan, S. Q. Shen, B. Wang, Q. Dai, S. J. Altschuler, and L. F. Wu. Integrative spatial analysis of cell morphologies and transcriptional states with muse. Nature biotechnology, 40(8):1200–1209, 2022.

7. Y. Bengio, A. Courville, and P. Vincent. Representation learning: A review and new perspectives. IEEE transactions on pattern analysis and machine intelligence, 35(8):1798–1828, 2013.

8. V. D. Blondel, J.-L. Guillaume, R. Lambiotte, and E. Lefebvre. Fast unfolding of communities in large networks. Journal of statistical mechanics: theory and experiment, 2008(10):P10008, 2008.

9. A. Borji. Pros and cons of gan evaluation measures. Computer vision and image understanding, 179:41–65, 2019.

10. T. Chen, S. Kornblith, M. Norouzi, and G. Hinton. A simple framework for contrastive learning of visual representations. In International conference on machine learning, pages 1597–1607. PMLR, 2020.

11. D. T. Forster, S. C. Li, Y. Yashiroda, M. Yoshimura, Z. Li, L. A. V. Isuhuaylas, K. Itto-Nakama, D. Yamanaka, Y. Ohya, H. Osada, et al. Bionic: biological network integration using convolutions. Nature Methods, pages 1–12, 2022.

12. R. Girdhar, A. El-Nouby, Z. Liu, M. Singh, K. V. Alwala, A. Joulin, and I. Misra. Imagebind: One embedding space to bind them all. In Proceedings of the IEEE/CVF Conference on Computer Vision and Pattern Recognition, pages 15180–15190, 2023.

13. M. Giurgiu, J. Reinhard, B. Brauner, I. Dunger-Kaltenbach, G. Fobo, G. Frishman, C. Montrone, and A. Ruepp. Corum: the comprehensive resource of mammalian protein complexes—2019. Nucleic acids research, 47(D1):D559–D563, 2019.

14. X. Glorot and Y. Bengio. Understanding the difficulty of training deep feedforward neural networks. In Proceedings of the thirteenth international conference on artificial intelligence and statistics, pages 249–256. JMLR Workshop and Conference Proceedings, 2010.

15. I. Goodfellow, J. Pouget-Abadie, M. Mirza, B. Xu, D. Warde-Farley, S. Ozair, A. Courville, and Y. Bengio. Generative adversarial nets. Advances in neural information processing systems, 27, 2014.

16. A. Grover and J. Leskovec. node2vec: Scalable feature learning for networks. In Proceedings of the 22nd ACM SIGKDD International Conference on Knowledge Discovery and Data Mining, KDD ’16, pages 855–864, New York, NY, USA, Aug. 2016. Association for Computing Machinery.

17. I. Gulrajani, F. Ahmed, M. Arjovsky, V. Dumoulin, and A. C. Courville. Improved training of wasserstein gans. Advances in neural information processing systems, 30, 2017.

18. M. Heusel, H. Ramsauer, T. Unterthiner, B. Nessler, and S. Hochreiter. Gans trained by a two time-scale update rule converge to a local nash equilibrium. Advances in neural information processing systems, 30, 2017.

19. E. L. Huttlin, R. J. Bruckner, J. Navarrete-Perea, J. R. Cannon, K. Baltier, F. Gebreab, M. P. Gygi, A. Thornock, G. Zarraga, S. Tam, et al. Dual proteome-scale networks reveal cell-specific remodeling of the human interactome. Cell, 184(11):3022–3040, 2021.

20. J. Jumper, R. Evans, A. Pritzel, T. Green, M. Figurnov, O. Ronneberger, K. Tunyasuvunakool, R. Bates, A. Žídek, A. Potapenko, et al. Highly accurate protein structure prediction with alphafold. nature, 596(7873):583–589, 2021.

21. M. Kanehisa and S. Goto. KEGG: kyoto encyclopedia of genes and genomes. Nucleic Acids Res., 28(1):27–30, Jan. 2000.

22. D. P. Kingma and J. Ba. Adam: A method for stochastic optimization. arXiv preprint arXiv:1412.6980, 2014.

23. L. Kondratyeva, I. Alekseenko, I. Chernov, and E. Sverdlov. Data incompleteness may form a hard-to-overcome barrier to decoding life’s mechanism. Biology, 11(8):1208, 2022.

24. D. Lahat, T. Adali, and C. Jutten. Multimodal data fusion: an overview of methods, challenges, and prospects. Proceedings of the IEEE, 103(9):1449–1477, 2015.

25. Z. Lin, H. Akin, R. Rao, B. Hie, Z. Zhu, W. Lu, N. Smetanin, R. Verkuil, O. Kabeli, Y. Shmueli, et al. Evolutionary-scale prediction of atomic-level protein structure with a language model. Science, 379(6637):1123–1130, 2023.

26. X. Mao, Q. Li, H. Xie, R. Y. Lau, Z. Wang, and S. Paul Smolley. Least squares generative adversarial networks. In Proceedings of the IEEE international conference on computer vision, pages 2794–2802, 2017.

27. L. McInnes, J. Healy, and J. Melville. Umap: Uniform manifold approximation and projection for dimension reduction. arXiv preprint arXiv:1802.03426, 2018.

28. R. Nasser, L. V. Schaffer, T. Ideker, and R. Sharan. Multi-modal contrastive learning of subcellular organization using dice. Bioinformatics, 40(Supplement 2):ii105–ii110, 2024.

29. J. Ngiam, A. Khosla, M. Kim, J. Nam, H. Lee, and A. Y. Ng. Multimodal deep learning. In Proceedings of the 28th international conference on machine learning (ICML-11), pages 689–696, 2011.

30. W. Ouyang, C. F. Winsnes, M. Hjelmare, A. J. Cesnik, L. Åkesson, H. Xu, D. P. Sullivan, S. Dai, J. Lan, P. Jinmo, et al. Analysis of the human protein atlas image classification competition. Nature methods, 16(12):1254–1261, 2019.

31. A. Radford. Unsupervised representation learning with deep convolutional generative adversarial networks. arXiv preprint arXiv:1511.06434, 2015.

32. P. J. Thul, L. Åkesson, M. Wiking, D. Mahdessian, A. Geladaki, H. Ait Blal, T. Alm, A. Asplund, L. Björk, L. M. Breckels, et al. A subcellular map of the human proteome. Science, 356(6340):eaal3321, 2017.

33. T. Wang and P. Isola. Understanding contrastive representation learning through alignment and uniformity on the hypersphere. In International Conference on Machine Learning, pages 9929–9939. PMLR, 2020.

34. F. Zheng, S. Zhang, C. Churas, D. Pratt, I. Bahar, and T. Ideker. Hidef: identifying persistent structures in multiscale ‘omics data. Genome biology, 22(1):1–15, 2021.

35. J.-Y. Zhu, T. Park, P. Isola, and A. A. Efros. Unpaired image-to-image translation using cycle-consistent adversarial networks. In Proceedings of the IEEE international conference on computer vision, pages 2223–2232, 2017.

